# A novel time-lapse imaging method for studying developing bacterial biofilms

**DOI:** 10.1101/2022.06.24.497465

**Authors:** Momir Futo, Tin Široki, Sara Koska, Nina Čorak, Anja Tušar, Mirjana Domazet-Lošo, Tomislav Domazet-Lošo

## Abstract

In nature, bacteria prevailingly reside in the form of biofilms. These elaborately organized surface-bound assemblages of bacterial cells show numerous features of multicellular organization. We recently showed that biofilm growth is a true developmental process, which resembles developmental processes in multicellular eukaryotes. To study the biofilm growth, in a fashion of eukaryotic ontogeny, it is essential to define dynamics and critical transitional phases of this process. The first step in this endeavor is to record the gross morphological changes of biofilm ontogeny under standardized conditions. This visual information is instrumental in guiding the sampling strategy for the later omics analyses of biofilm ontogeny. However, none of the currently available visualizations methods is specifically tailored for recording gross morphology across the whole biofilm development. To address this void, here we present an affordable Arduino-based approach for time-lapse visualization of complete biofilm ontogeny. The major challenge in recording biofilm development on the air-solid interphase is water condensation, which compromises filming directly through the lid of a Petri dish. To overcome these trade-offs, we developed an Arduino microcontroller setup which synchronizes a robotic arm, responsible for opening and closing the Petri dish lid, with the activity of a stereomicroscope-mounted camera and lighting conditions. We placed this setup into microbiological incubator that maintains temperature and humidity during the biofilm growth. As a proof-of-principle, we recorded biofilm development of five *Bacillus subtilis* strains that show different morphological and developmental dynamics.

## Introduction

Bacterial biofilms are the most common life-form on Earth^1^. They are highly organized surface-associated communities of bacterial cells embedded in a self-derived extracellular matrix^2^. From the origin of life, bacterial biofilms are continuously present in the most diverse habitats where they show stunning ecological adaptations^3,4^. Due to their ecological, medical^5,6^ and commercial importance^7^, bacterial biofilms have been studied from many angles using state-of-the-art omics approaches^8–10^. As an example, we recently showed that biofilm growth represents a true developmental process that contains discrete developmental phases comparable to those of developing eukaryotic embryos^10^.

To achieve these findings, we transferred experimental and analytical approaches regularly used in developmental biology of multicellular eukaryotes to the analysis of *Bacillus subtilis* biofilm ontogeny^10–13^. Traditionally, the study of eukaryotic development starts with a careful description of gross morphological changes along the ontogeny; from fertilization until the formation of an adult organism. To ensure reproducibility and allow comparisons across studies, the developmental process of a eukaryotic organism is usually divided into stages that are temporally and morphologically defined under standardized laboratory conditions^12^. However, these developmental stages not only help researchers to navigate along the developmental process, but also reflect underlying molecular processes that are organized in a discrete fashion^10,12,14–16^. It is, therefore, important that the gross morphological changes of biofilm ontogeny are recorded and defined before the biofilm sampling for various omics analyses is performed.

To optimize sampling strategy for downstream transcriptomic and proteomic analysis, we previously made a time-lapse video of complete *B. subtilis* development on a solid-air interface^10^. However, during these experiments we faced substantial methodological challenges during the recording of time-lapse videos related to the obstruction of visual clarity due to the water condensation on Petri dish lids (Fig. 1). The same condensation-induced photo blurriness was noticed previously in a comparable effort to record bacterial colony growth by time-lapse visualization^17^. An obvious solution to this problem would be to entirely remove the lid during the filming process. However, this approach compromises the experiment by increasing desiccation and contamination of the agar medium in a Petri dish. To address these tradeoffs, in our previous work we applied a brute-force approach by manually opening, filming and subsequently closing the Petri dish lid during the experiment^10^. However, this solution is rather impractical, or even prohibitive, as it required engagement of substantial workforce over the weeks-long filming sessions of biofilm development, which required shooting in 15 min intervals. Another drawback associated with this solution is that frequent opening and closing of the incubator door leads to temperature and humidity fluctuations, which in turn must be carefully controlled. In addition, the same procedure increases the probability of airborne contaminants in the incubator space.

**Figure 1.**
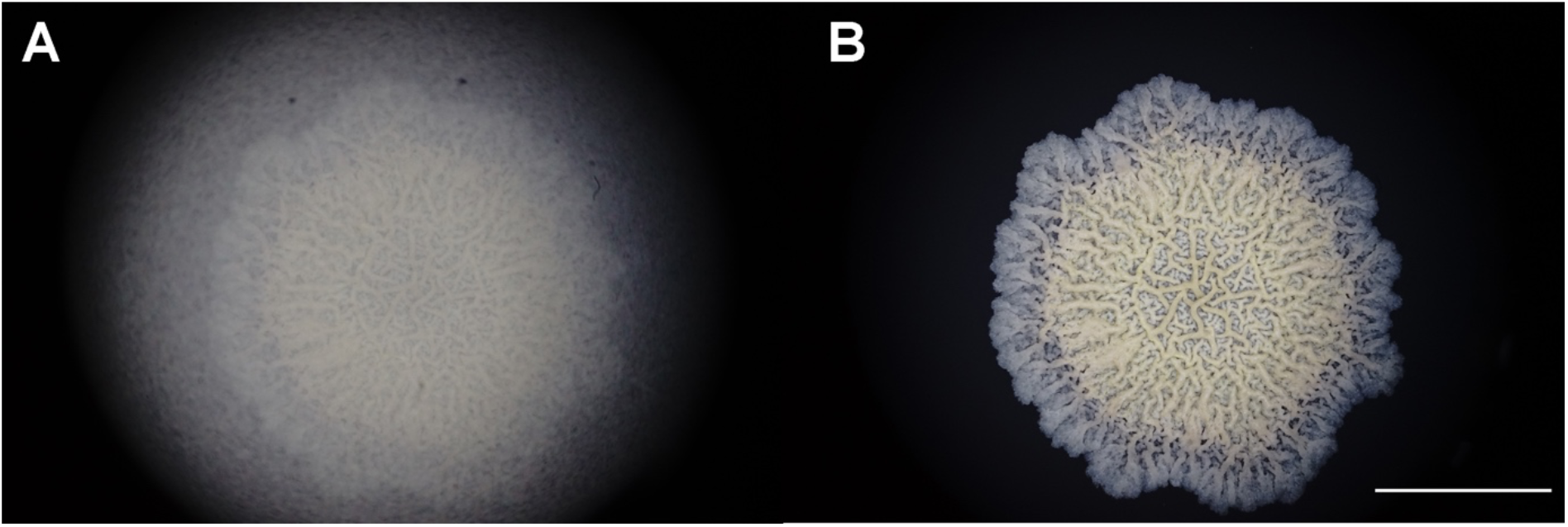
Water condensation on the Petri dish lid obstructs visual clarity during time-lapse visualization of *Bacillus subtilis* biofilm development. *Bacillus subtilis* NCIB 3610 biofilm photographed through a Zeiss Stemi C-2000 stereomicroscope at a 0.8x magnification on agar media in a Petri dish. The scale bar indicates 1cm. | **A** – Photo taken through the lid of a Petri dish obstructed by water condensation, **B** – The photo of the same biofilm taken with the removed lid of the Petri dish.

To address these issues, we here present an Arduino microcontroller-based solution for time-lapse visualization of developing biofilms at solid-air interfaces. As a proof of concept, we recorded time-lapse videos of five *B. subtilis* strains developing biofilms that show distinct biofilm colony morphologies and developmental dynamics.

## Results and Discussion

### Technical aspects of the method

In a closed incubator environment, the standard growth temperature for mesophilic bacteria, which cover the majority of common environmental bacteria and human pathogens, ranges between 20 and 45 °C^18^. This relatively high incubator temperature causes an extensive water evaporation from agarose-based growth media, which often directly affects the growth of bacterial colonies. During standard bacterial incubations this excess evaporation is prevented by a Petri dish lid. However, slight temperature differences between the lid and the inner Petri dish space often lead to moisture condensation on the inner Petri dish lid surface^19^. In the majority of microbiological experiments this phenomenon is of no concern, but in biofilm macrocolony photography and the production of time-lapse videos, the vapor condensation on the transparent lid causes considerable difficulties in taking clear biofilm photographs (Fig. 1). This is clearly visible in some time-lapse videos of a previous methodological paper that deals with bacterial macrocolony photography^17^.

To overcome these obstacles, we constructed an Arduino-controlled biofilm visualization setup (Fig. 2). The core of this setup, which consists of a camera mounted on top of a stereomicroscope (Fig. 2E) and an Arduino microcontroller with robotic arm (Fig. 2F), is placed inside a microbiological incubator (Fig. 2A). The basic logic behind this apparatus is to automatize the Petri dish lid opening, synchronize this step with the automated camera shots of a growing biofilm, and perform these actions under tightly controlled temperature, humidity and lightning conditions. The main functional part of our setup is a small, acrylic robotic arm controlled by an Arduino microcontroller (Fig. 2F). The sole purpose of this robotic arm, which is firmly attached to the Petri dish lid, is to open and close the Petri dish in short, preprogramed photo-intervals that allow visually unobstructed shooting of a biofilm that grows on an agar plate (Fig. 3).

**Figure 2.**
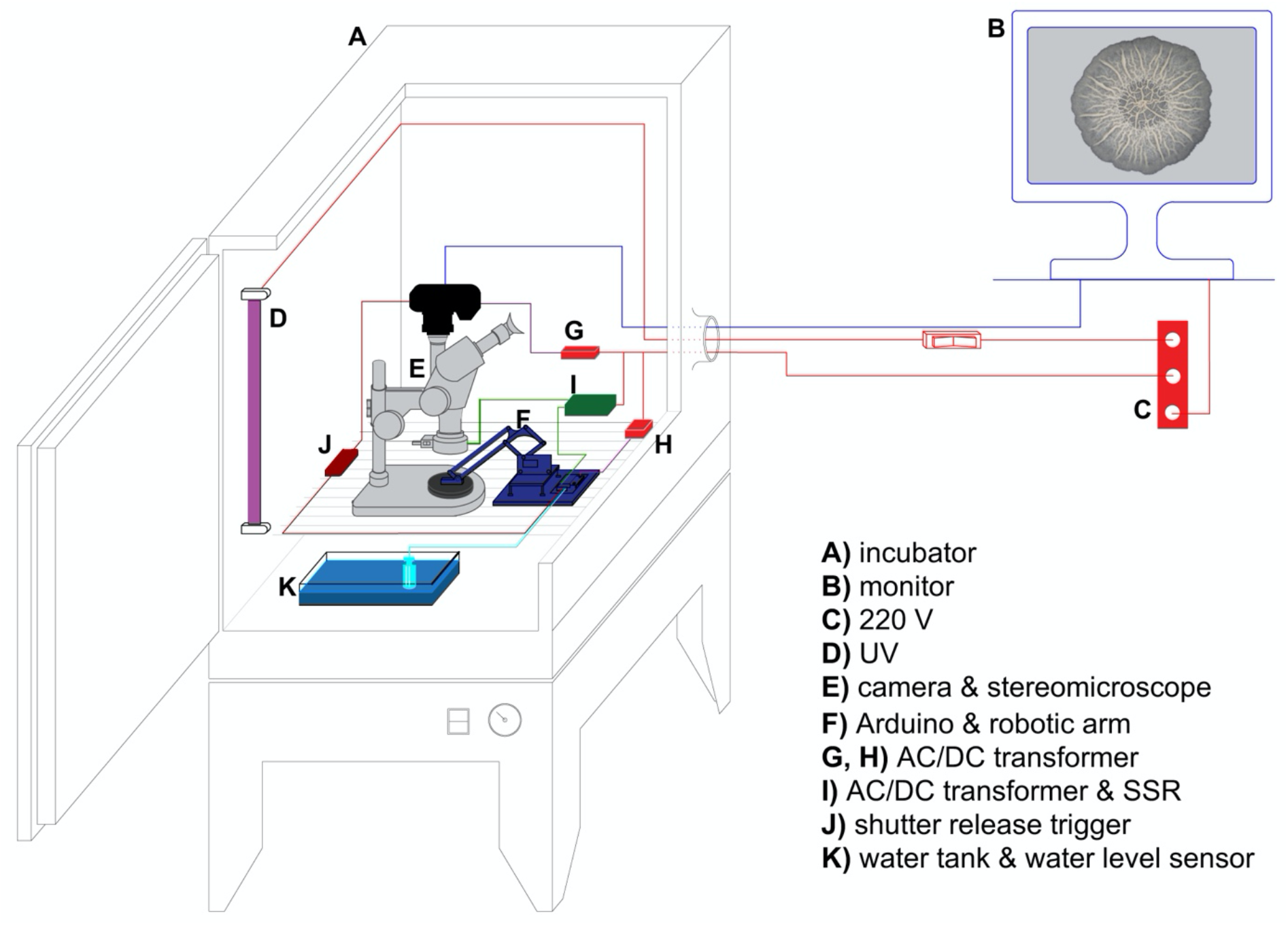
The experimental apparatus setup. **A** – microbiological incubator, **B** – an external computer monitor attached to the camera, **C** – 220V AC outlet, **D** – UV lamp, **E** – mirrorless SONY alpha 7 II camera coupled with a 144 LED light-ring mounted on a Zeiss Stemi C-2000 stereo microscope using a T2 adapter, **F** – Arduino microcontroller with an acrylic robotic arm, **G, H** – AC/DC transformers, **I** – AC/DC transformer and a solid state relay module (Omron), **J** – digital timer remote MC-36B (Neewer), **K** – water-filled container with a water level detection sensor module.

**Figure 3.**
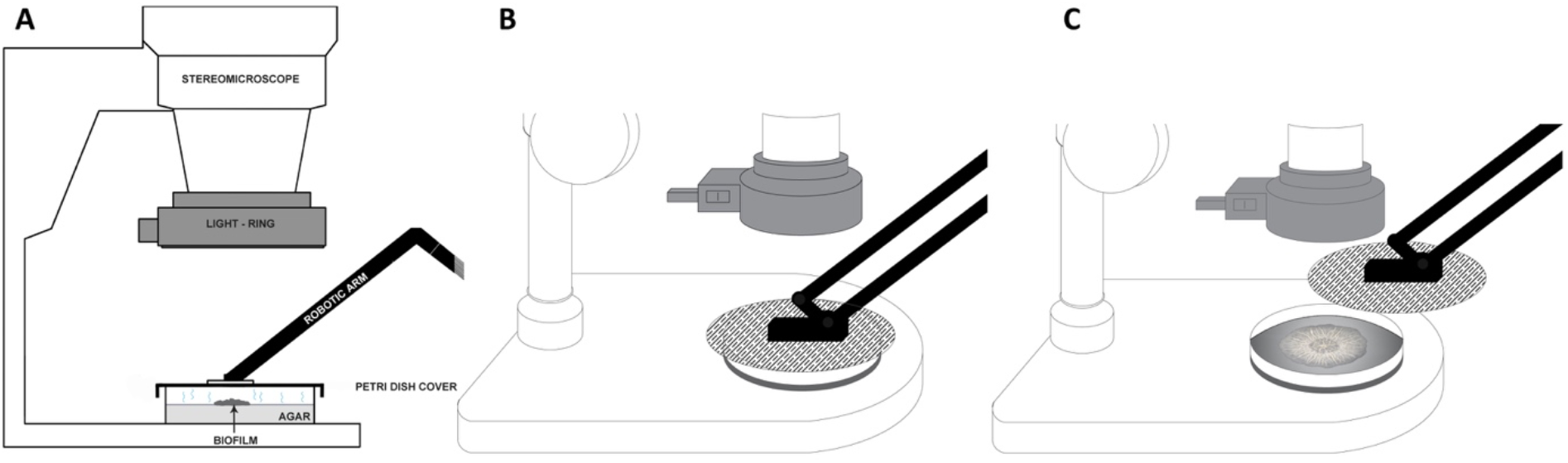
The principle of the robotic arm actions. **A** – sideview of the closed agar plate with a developing biofilm, **B** – the closed Petri dish with a developing biofilm between shootings, **C** – the robotic arm opens the Petri dish before the camera shooting will take place.

During the whole experiment the robotic arm is firmly attached to a Petri dish lid. In the resting periods — *i*.*e*. before the start of the experiment, after the last photo has been taken, and between the two recording steps — the robotic arm keeps the lid in a closed Petri dish position (Fig. 3B), thus preventing the unnecessary medium desiccation and contamination. The Arduino microcontroller starts the time-lapse recording sequence by switching on the LED light-ring mounted on the objective of the stereomicroscope (Fig. 2E, 4F) via the solid-state relay module (Fig. 4E).

**Figure 4.**
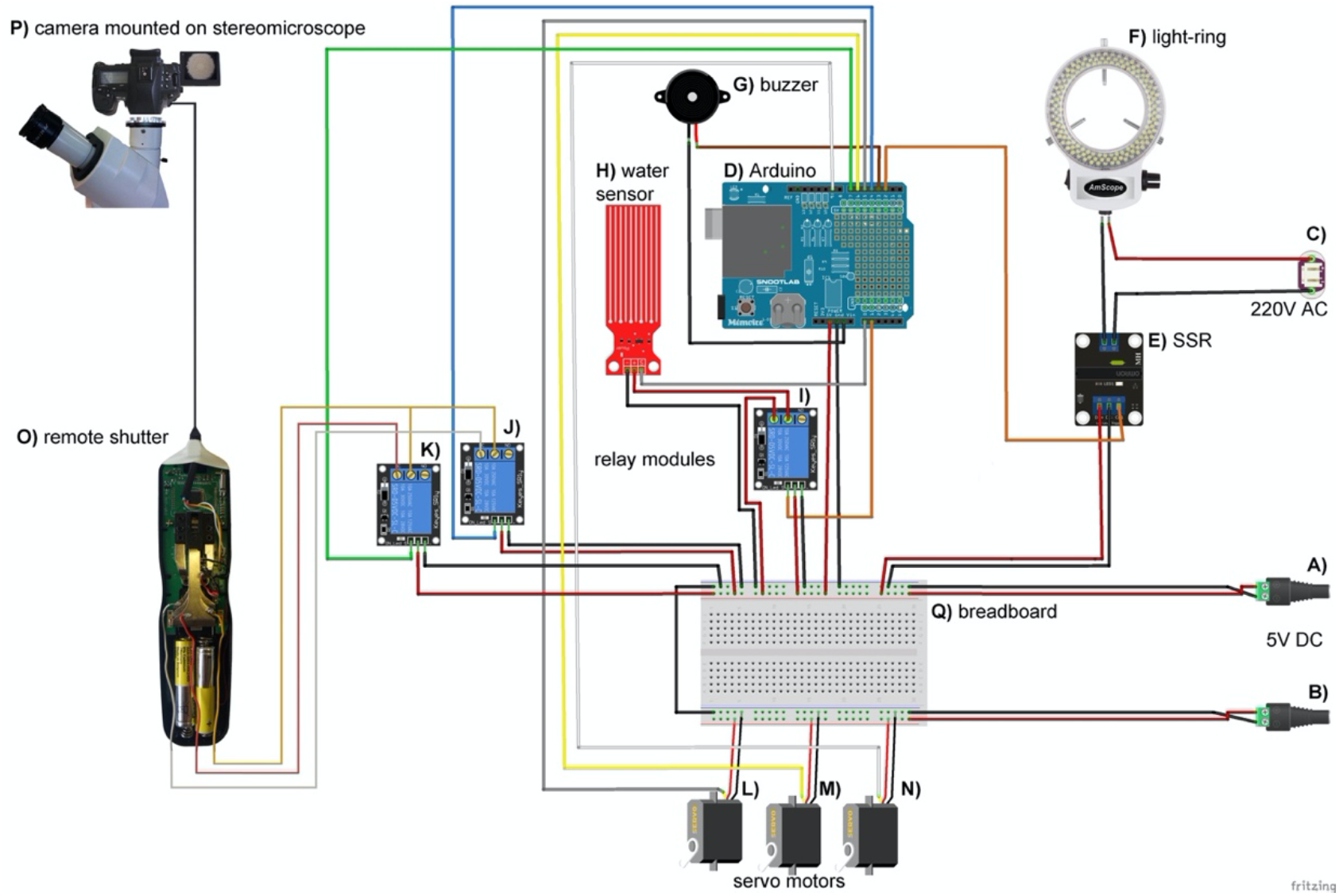
Arduino microcontroller wiring schematics. **A, B** – 5V DC power sources, **C** – 220V AC power source, **D –** Arduino® Uno R3 microcontroller board, **E** – solid state relay module (Omron), **F** – 144 LED light-ring for stereo microscopes (AmScope), **G** - passive buzzer, **H** – water level detection sensor module, **I, J, K** – 5V DC Arduino KY-019 relay modules, **L, M, N** – SG90 servo motors, **O** – digital timer remote shutter release trigger MC-36B (Neewer), **P** – SONY alpha 7 II mirrorless camera mounted on a Zeiss Stemi C-2000 stereo microscope using a T2 adapter, **Q** – 30-row solderless breadboard with two bus stripes. Arduino schematic was developed using the Fritzing software V0.9.3^20^ and Adobe Photoshop CC 2017.

Several milliseconds later, the robotic arm attached to the lid first moves several centimeters vertically, and then performs a side move to get the lid out of the camera’s field of view (Fig. 3C). During these movements the lid keeps a parallel position to the Petri dish that contains a growing biofilm. At the moment when the robotic arm stops in the opened Petri dish position (Fig. 3C), the microcontroller takes a photo of the biofilm via the remote shutter (Fig. 2J, 4O) connected to the camera (Fig. 2E, 4P). After the photo has been recorded, the Petri dish is closed again by the same robotic arm movements, but in the opposite direction. Several milliseconds after the closed Petri dish position is reached, the Arduino microcontroller switches off the light-ring. The next cycle of time-lapse recording is repeated depending on the time intervals entered in the software that drives the microcontroller.

To further minimize the dehydration of agar media during the filming of biofilm growth, we placed four square-shaped plastic containers filled with distilled water on the bottom of the incubator covering approximately 78% of the bottom area. These water-filled containers were performing the role of a humidifier, which allowed us to maintain relative air humidity within the incubator at approximately 85% during the visualization experiments that lasted up to 21 days. In the case a longer experimental growth period would be needed, our setup can be easily upgraded with a double-decker agar hydration design described in Peñil Cobo et al. 2018. To control the water level in water containers, we connected a water-level sensor to the Arduino microcontroller (Fig. 2K, 4H). In situations when the water level dropped below the critical threshold and the water containers needed refilling, the Arduino microcontroller sounded an alarm over the passive buzzer (Fig. 4G). We prevented unnecessary electrode corrosion of the water-level sensor through electrolysis by connecting it through a relay module (Fig. 4I), which switched on the sensor only during the measurement time.

In order to detect an occasional camera focus loss due to biofilm growth in three dimensions and to prevent eventual mechanical misbehaviors of the system, it is necessary to visually control the activity of the setup during the filming session. However, to avoid the frequent opening of the incubator door, which in turn leads to humidity, lighting and temperature fluctuations within the incubator chamber, we connected an external monitor to the camera that allows real-time monitoring of biofilm growth and setup behavior (Fig. 2B).

Before the start of our visualization experiments, we sterilized the inner parts of the incubator, its atmosphere and the whole apparatus by a UV sterilizer that we added in the incubator chamber (Fig. 2D). This procedure allowed us to largely prevent contamination of agar media during the experiments. Nevertheless, we had to occasionally open the incubator during the experiment to manually correct the focus of the stereomicroscope that became suboptimal due to changes in the biofilm 3D morphology, or to refill the water containers. In the vast majority of experiments this was not a problem, but occasionally a spore contamination was unavoidable. However, from our experience, the air-borne fungal contamination was not visible on the agar-plates before two weeks of continuous recording (Supplementary Video 5). In order to further minimize the fungal or bacterial spore contamination of the agar-plates, an automated stereomicroscope focusing mechanism, as well as an automated water container refill mechanism, should be added to the setup in the future.

### Proof of concept: Strain-specific development in Bacillus subtilis

To demonstrate our time-lapse visualization method, we recorded developmental and morphological changes during the biofilm growth of five distinctive *B. subtilis* strains (Fig. 5).

**Figure 5.**
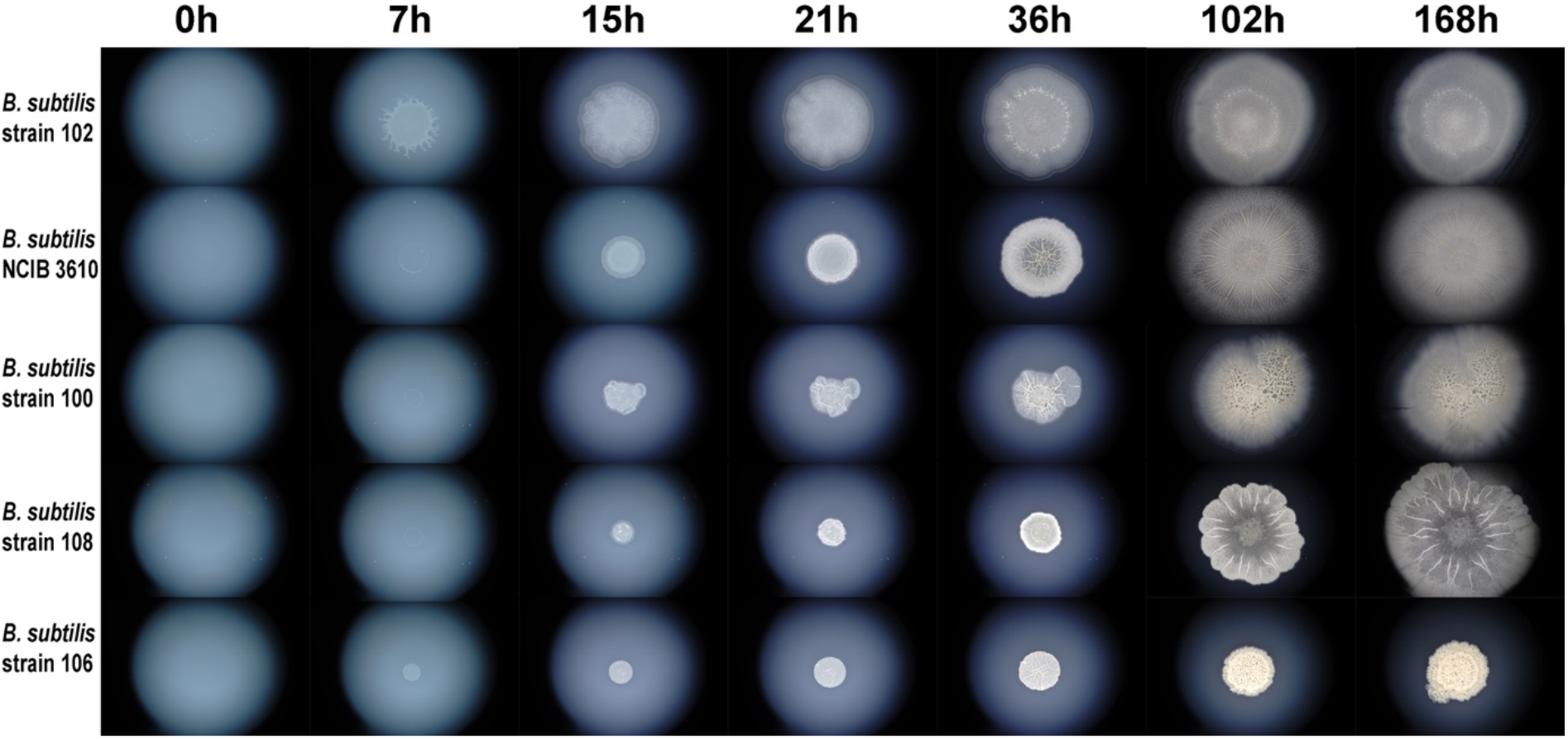
Selected time-lapse photos of developing *B. subtilis* biofilms. We recorded the biofilm development of five *B. subtilis* strains (102, NCIB3610, 100, 108, 106) using our time-lapse setup. Corresponding photos in seven time points are shown (0h, 7h, 15h, 21h, 36h, 102h, 168h). The full time-lapse recordings for every strain that covers seven day period are available in Supplementary Video 1, 2, 3, 4 & 5. These time-lapse recordings reveal that five *B. subtilis* strains show a remarkable difference in developmental dynamics and biofilm morphology.

We photographed biofilm development of every strain independently in photo-intervals of 6 min. This recording frequency allowed us to produce detailed time-lapse videos with an optimal frame rate (30 fps). We recorded 1,681 photos for every of the four *B. subtilis* strains (102, 100, NCIB 3610, and 108) that we grew for seven days. In addition, we collected 5,041 photos of a slowly developing strain (106) that we grew for 21 days. To give an overview of the recording process, we show seven representative photos recorded during seven-day period for all five strains (Fig. 5). However, as the full dynamics of biofilm growth and the changes in its morphology are only possible to grasp from time-lapse videos, we also assembled our photo collections into five time-lapse videos that show the biofilm growth of every strain independently (Supplementary Video 1, 2, 3, 4 & 5). For comparative purposes, we also compiled two additional videos that show the growth of two strains (102, 106) and four strains (102, 100, NCIB3610 and 108) in parallel (Supplementary Video 6 & 7).

The first obvious difference among the observed *B. subtils* strains was their growth speed (Fig. 5, Supplementary Video 6). Although all observed strains grew under the same standardized environmental conditions (1.5% w/v agarose, 30 °C and 85% relative humidity), they showed rather different growth dynamics (Fig. 5, Supplementary Video 6 & 7). For instance, the 102 strain showed the fastest biofilm expansion, in contrast to the 106 strain which showed the slowest growth within the seven-day timeframe (Fig. 5, Supplementary Video 7). Except the fastest growth, the 102 strain showed a star-shaped morphology in early phase (7h) of biofilm growth that was absent in other strains (Fig. 5, Supplementary Video 2, 6, 7).

Besides the growth rate, in our time-lapse videos one could also look at the changes in colony morphology, which is considered one of the hallmarks of *B. subtilis* biofilms^21^. In our previous work we showed that typical wrinkled biofilm morphology in the NCIB 3610 strain starts to develop in the transition stage during the second day of biofilm growth, which correlates with dramatic changes in expression patterns^10^. In this study we were able to reproduce this pattern in the same strain using our automated apparatus recording. The time-lapse video shows that wrinkles first start to appear in the centre of NCIB 3610 biofilm at the onset of the second day of biofilm growth (Supplementary Video 4). In comparison to the NCIB 3610 strain, the wrinkles in other four strains showed different morphology in the terms of density, timing of their first appearance, and three-dimensional structure (Fig. 5, Supplementary Video 6 & 7). This developmental variability between strains suggests differences in underlying expression patterns that could be further explored by time resolved sampling and subsequent transcriptomic and proteomic analysis^10^.

## Conclusions

We developed a novel, automated time-lapse imaging method for studying the gross morphology of developing bacterial biofilms over long periods of time that solves the problem of condensation-induced photo blurriness. As our method is based on an easily programmable Arduino microcontroller platform, it is highly flexible in terms of controlling photo periods for time-lapse photography. In addition, our setup could be easily upgraded with various additional elements such as diverse kinds of sensors that might come useful in different experimental setups. Although our visualization setup is primarily developed for visualization of biofilms or macrocolonies that grow on solid-air interfaces, with minimal modifications it could be also used for visualization of biofilms that float on liquid-air interfaces as pellicles. Due to the relatively low cost of its main components, this method is an affordable solution for microbiological time-lapse recording in various scientific and educational environments. The quality of the time-lapse videos that could be produced by this setup was already acknowledged with an Honorable mention award at the 2021 Nikon Small World in Motion Competition^22^.

## Material and methods

### Bacterial strains

#### Isolation, identification and storage

For the purpose of time-lapse visualization, we used five strains of the bacterial model species *Bacillus subtilis*. A widely-used, biofilm-forming *B. subtilis* subsp. *subtilis* str. NCIB 3610 (strain 3610) was obtained from the Bacillus Genetic Stock Center (BGSC, Ohio State University, Columbus, OH, USA) and stored in 25% glycerol stocks at -80 °C. Four additional *B. subtilis* environmental strains were isolated from a topsoil sample collected from Ruđer Bošković Institute backyard following the below-described procedure. The five grams of topsoil were mixed with 45 mL of re-distilled, filter-sterilized water and vortexed for 3 min in a 50 mL falcon tube at room temperature (RT). After the soil precipitate has sedimented for 10 min at RT, the supernatant was serially diluted in steps of 10^−1^, 10^−2^ and 10^−3^. 100 μL of the undiluted and three diluted supernatants were plated out on 90 mm Petri dishes containing biofilm-promoting MSgg agar (5 mM potassium phosphate pH 7, 100 mM MOPS pH 7, 2 mM MgCl_2_, 700 μM CaCl_2_, 50 μM MnCl_2_, 50 μM FeCl_3_, 1 μM ZnCl_2_, 2 μM thiamine, 0.5% glycerol, 0.5% glutamate, 50 μg/mL tryptophan, 50 μg/mL phenylalanine solidified with 1.5% agar) using a Drigalski spatula in 10 replicates each. In total 40 inoculated MSgg plates were incubated for 24h at 30 °C. After incubation, all plates were visually inspected using a Stemi C-2000 stereo microscope (Zeiss). In total, 14 single-standing colonies that showed a visually interesting and attractive colony structure were isolated and inoculated on a fresh MSgg plates, each in three replicates. After 24h of incubation at 30 °C, a single colony from each of the 14 samples was analyzed using the MALDI-TOF Biotyper mass spectrometry system (Bruker Daltonik). Four out of 14 samples with the best and second-best match values above 2.0 to *B. subtilis* were chosen for the downstream visualization experiments as *B. subtilis* strains. These isolated strains were arbitrary named as strain 100, 102, 106, and 108 and were stored in 25% glycerol stocks at -80 °C.

#### Growth conditions before and during the visualization experiments

Two days before each visualization experiment, the bacteria from the -80 °C glycerol stock were plated on a LB agar plate (1% Bacto tryptone, 0.5% Bacto yeast extract, 1% NaCl, 1 mM NaOH solidified with 1.5% agar) and incubated for 24h at 30 °C. 10 mL of liquid LB medium were inoculated with a single colony and incubated overnight (16h) in a shaker at 30 °C and 250 rpm. On the day of the experiment, the plastic bottom of a 90 mm Petri dish containing MSgg agar was evenly painted with black acrylic paint (Marbau GmbH & Co. KG) and left upside down on the bench to dry on RT. After the paint had dried, the Petri dish was placed in the incubator, centered under the stereomicroscope objective (Zeiss Stemi C-2000, Fig. 2E, 3A) and glued to the stereomicroscope’s stage plate with three drops of silicone glue. The incubator was closed and sterilized with a 254 nm OFR UV light for 30 min (Fig. 2D). Following the incubator surface-sterilization, the Petri dish lid was removed and the MSgg agar plate was inoculated with 5 μL of the bacterial overnight culture while observed through the monitor attached to the camera mounted on the stereomicroscope (Fig. 2B), The 5 μL drop was pipetted in the middle of the field of view and focused using the stereomicroscope’s focus nob. Subsequently, the inoculated plate was covered with Petri dish cover attached to the robotic arm (Fig. 3B). The experiment started by closing the incubator, and turning the Arduino microcontroller ON. For the purpose of the five visualization experiments described in this study, biofilms were grown at 30°C, for 7 days (strains 100, 102, 108 and NCIB 3610) and 21 days (strain 106) respectively, while the relative air humidity inside the incubator was maintained at around 85%.

### Arduino controlled time-lapse photography setup

#### The robotic arm

The bacterial biofilm growth visualization of the five *B. subtilis* strains over various time-courses (7 and 21 days) were performed using the time-lapse stereomicroscope-photography apparatus shown in Fig. 2 and 4. The whole setup was operated by an Arduino Uno R3 microcontroller board (Arduino) (Fig. 4D). Digital pins 5, 6 and 9 of the Arduino were each assigned to a SG90 servomotor (Fig. 4L, M, N), respectively. The servomotors were moving a simple, acrylic robotic arm in three axes. The robotic arm was attached to the lid of Petri dish with an adhesive tape. All three servomotors were powered by a DC 5V power-source (Fig. 4B) running separately from the main DC 5V line that powered the microcontroller and the modules (Fig. 4A).

#### Light, temperature and humidity

Optimal brightness for successful photography in the dark incubator was provided by an episcopic 144 LED light-ring (Fig. 4F) mounted around the stereomicroscope’s objective (Fig. 2E, 3). The periodical and preprogramed ON/OFF switching of the light-ring was governed by the microcontroller-mediated solid-state relay module (Omron) (Fig. 4E), which was closing and opening the light-ring’s DC 12V circuit provided by the DC 12V power-source (Fig. 4C). The relay was connected to the microcontroller via digital pin 2.

The optimal temperature for bacterial growth was provided by the microbiological incubator (Termomedicinski aparati – TMA) (Fig. 2A). All bacterial strains were grown at 30 °C. The air-humidity in the incubator was maintained by evaporation from four water containers placed in the incubator and filled with redistilled water (Fig. 2K). The water level in the containers was controlled with a water level sensor detection module (Fig. 4H). The sensor was connected to the microcontroller’s analog input pin A0. The 5V DC current powering the sensor was periodically switched ON/OFF by the 5V DC KY-019 relay module (Fig. 4I). The relay module was connected to the analog input pin A1 of the microcontroller. Digital pin 3 was connected to a passive buzzer (Fig. 4G). The humidity was monitored with a HM16 thermo/hygrometer (Bauer GmbH) placed within the incubator.

#### The photography and camera settings

The photographs of growing biofilms were recorded using a SONY alpha 7 II mirrorless camera (Fig. 4P) mounted on a Stemi C-2000 (Zeiss) stereo microscope using a T2 adapter (Fig. 2E). Digital pins 4 and 7 of the microcontroller board were connected to two DC 5V KY-019 relay modules (Fig. 4J, K), respectively. Once triggered by a signal received from the microcontroller, the two relay modules (Fig. 4K, L) would successively, with a 1s time-gap, close the two 1.5V DC circuits of the digital timer remote shutter release trigger MC-36B (Neewer) (Fig. 2J, 4O). Such an action would imitate a physical finger-push on the shutter release trigger and as a result would produce and send a signal to the camera connected to the shutter release trigger via a USB connection. The photographs were taken in six min intervals during seven days (strain 100, 102, 108 and NCIB 3610) and 21 days (strain 106), respectively. The magnification of the stereomicroscope was set to 0.8 x. The camera was set to the “Aperture Priority” mode with the following parameters: image size: 24M; image quality: RAW+J; image resolution: 6000×4000 px, 350 dpi; drive mode: single shooting; flash compensation: ±0.0; focus area: center; exposure compensation: ±0.0; light sensitivity ISO: 50; metering mode: multi; white balance: AWB; creative style – “Clear” while the D-Range Optimizer and flash were set to “off”.

### Production of time-lapse videos

The time-lapse videos of developing biofilm macrocolonies were produced from series of JPEG photographs using the Adobe After Effects CC 2017 software. Time-lapse videos of *B. subtilis* strains 102, 100, NCIB 3610 and 108 biofilms were assembled of four different photography time-series sets, each containing 1,681 photographs. The time-lapse video of the strain 106 was rendered using 5,041 photographs of the developing biofilm. The render settings were set to: “Best”, video size was 1920 × 1080 px in full resolution, auto input was switched off. The frame rate in all videos was set to 30 fps, the format of the videos was QuickTime, while the codec used was MPEG-4 Video.

## Acknowledgements

We thank Agneš Beneš for her help in graphical design, Tomislav Mrla and Kristian Vlahoviček for their help with electronics, Mirjana Filipović for the technical support, and Tomislav Ivanković for the fruitful comments and discussions. This study was supported by the Croatian Science Foundation under the project IP-2016-06-5924, the Adris Foundation, the European Regional Development Fund (KK.01.1.1.01.0009 DATACROSS), and the European Union’s Horizon 2020 research and innovation program under the Marie Sklodowska-Curie grant agreement 955626 (PEST-BIN).

## Author contributions

MF and TDL conceived the study; MF developed the visualization method; MF, SK, and AT performed the microbiological lab-work; MF, TŠ and MDL developed the microcontroller code; MF and TŠ edited the photographs; SK and AT tested the setup; MF and NČ produced the time-lapse videos; MF and NČ produced the figures; MF and TDL wrote the manuscript with contributions of all authors. All authors read and approved the final version of the manuscript.

## Data availability

The videos presented in this paper are available for immediate viewing on the following private YouTube links and for download on the following FigShare links (in brackets): Supplementary Video 1 – strain 100: https://youtu.be/t9phHaF3w4c (https://figshare.com/s/0437415630ba504d46fa); Supplementary Video 2 – strain 102: https://youtu.be/wVtxh-oSeXg (https://figshare.com/s/4b7befdb3b2342c11689); Supplementary Video 3 – strain 108: https://youtu.be/trrpci4uf0I (https://figshare.com/s/7ac0070ee0a85b92086c); Supplementary Video 4 – strain NCIB 3610: https://youtu.be/Kl6sDOqm6eo (https://figshare.com/s/358422b2cc948fb1d637); Supplementary Video 5 – strain 106: https://youtu.be/XZdVOzuULjs (https://figshare.com/s/b78c57d2466171d6b6a6); Supplementary Video 6 - strains 102, 100, NCIB 3610 and 108: https://youtu.be/CZ0CC9L6hdo (https://figshare.com/s/1748e92357823451685f) and Supplementary Video 7 - strains 102 and 106: https://youtu.be/qTX5QyFLyuA (https://figshare.com/s/b37f172c893c22d79f1d). The code driving the Arduino microcontroller and the Python script for photography time-tagging are available at the GitHub link https://github.com/bacillus-biofilms/time-lapse-imaging.

## Literature

1. Flemming, H.-C. & Wuertz, S. Bacteria and archaea on Earth and their abundance in biofilms. Nat. Rev. Microbiol. 17, 247–260 (2019). https://doi.org/10.1038/s41579-019-0158-9

2. Flemming, H.-C. & Wingender, J. The biofilm matrix. Nat. Rev. Microbiol. 8, 623–633 (2010). https://doi.org/10.1038/nrmicro2415

3. Yang, L. et al. Current understanding of multi-species biofilms. Int. J. Oral Sci. 3, 74–81 (2011). https://doi.org/10.4248/IJOS11027

4. Escudero, C., Vera, M., Oggerin, M. & Amils, R. Active microbial biofilms in deep poor porous continental subsurface rocks. Sci. Rep. 8, 1538 (2018). https://doi.org/10.1038/s41598-018-19903-z

5. Burmølle, M. et al. Biofilms in chronic infections – a matter of opportunity – monospecies biofilms in multispecies infections. FEMS Immunol. Med. Microbiol. 59, 324–336 (2010). https://doi.org/10.1111/j.1574-695X.2010.00714.x

6. Percival, S. L., Suleman, L., Vuotto, C. & Donelli, G. Healthcare-associated infections, medical devices and biofilms: risk, tolerance and control. J. Med. Microbiol. 64, 323–334 (2015). https://doi.org/10.1099/jmm.0.000032

7. Backer, R. et al. Plant Growth-Promoting Rhizobacteria: Context, Mechanisms of Action, and Roadmap to Commercialization of Biostimulants for Sustainable Agriculture. Front. Plant Sci. 9, 1473 (2018). https://doi.org/10.3389/fpls.2018.01473

8. Franklin, M. J., Chang, C., Akiyama, T. & Bothner, B. New Technologies for Studying Biofilms. Microbiol. Spectr. 3, 3.4.27 (2015). https://doi.org/10.1128/microbiolspec.MB-0016-2014

9. Azeredo, J. et al. Critical review on biofilm methods. Crit. Rev. Microbiol. 43, 313–351 (2017). https://doi.org/10.1080/1040841X.2016.1208146

10. Futo, M. et al. Embryo-Like Features in Developing Bacillus subtilis Biofilms. Mol. Biol. Evol. 38, 31–47 (2021). https://doi.org/10.1093/molbev/msaa217

11. Domazet-Lošo, T., Brajković, J. & Tautz, D. A phylostratigraphy approach to uncover the genomic history of major adaptations in metazoan lineages. Trends Genet. 23, 533–539 (2007). https://doi.org/10.1016/j.tig.2007.08.014

12. Domazet-Lošo, T. & Tautz, D. A phylogenetically based transcriptome age index mirrors ontogenetic divergence patterns. Nature 468, 815–818 (2010). https://doi.org/10.1038/nature09632

13. Shi, L. et al. Evolutionary Analysis of the Bacillus subtilis Genome Reveals New Genes Involved in Sporulation. Mol. Biol. Evol. 37, 1667–1678 (2020). https://doi.org/10.1093/molbev/msaa035

14. Monds, R. D. & O’Toole, G. A. The developmental model of microbial biofilms: ten years of a paradigm up for review. Trends Microbiol. 17, 73–87 (2009). https://doi.org/10.1016/j.tim.2008.11.001

15. Levin, M., Hashimshony, T., Wagner, F. & Yanai, I. Developmental Milestones Punctuate Gene Expression in the Caenorhabditis Embryo. Dev. Cell 22, 1101–1108 (2012). https://doi.org/10.1016/j.devcel.2012.04.004

16. Yanai, I. Development and Evolution through the Lens of Global Gene Regulation. Trends Genet. 34, 11–20 (2018). https://doi.org/10.1016/j.tig.2017.09.011

17. Peñil Cobo, M. et al. Visualizing Bacterial Colony Morphologies Using Time-Lapse Imaging Chamber MOCHA. J. Bacteriol. 200, (2018). https://doi.org/10.1128/JB.00413-17

18. Microbiology: a human perspective. (McGraw-Hill, 2001). ISBN-13: 978-0072318784

19. Finer, J. E. & Finer, J. J. A simple method for reducing moisture condensation on Petri dish lids. Plant Cell Tissue Organ Cult. 91, 299–304 (2007). https://doi.org/10.1007/s11240-007-9292-6

20. Knörig, A., Wettach, R. & Cohen, J. Fritzing: a tool for advancing electronic prototyping for designers. in Proceedings of the 3rd International Conference on Tangible and Embedded Interaction - TEI ‘09 351 (ACM Press, 2009). https://dl.acm.org/doi/10.1145/1517664.1517735

21. Branda, S. S., González-Pastor, J. E., Ben-Yehuda, S., Losick, R. & Kolter, R. Fruiting body formation by Bacillus subtilis. Proc. Natl. Acad. Sci. 98, 11621–11626 (2001). https://doi.org/10.1073/pnas.191384198

22. Futo, M. ‘5-day time-lapse of Bacillus subtilis biofilm growth and development’ Honorable Mention Award at the 2021 Nikon Small World in Motion Competition. (2021). https://www.nikonsmallworld.com/galleries/2021-small-world-in-motion-competition/time-lapse-of-bacillus-subtilis-biofilm-growth-and-development

